# Similar clinical improvement and maintenance after rTMS at 5 Hz using a simple vs. complex protocol in Alzheimer’s disease

**DOI:** 10.1101/232546

**Authors:** R. Alcalá-Lozano, E. Morelos-Santana, J.F. Cortés-Sotres, E. A. Garza-Villarreal, A. L. Sosa-Ortiz, J. J. Gonzalez-Olvera

## Abstract

**Brackground:** Current treatments for Alzheimer’s disease (AD) have a limited clinical response and methods, such as repetitive transcranial magnetic stimulation (rTMS), are being studied as possible treatments for the clinical symptoms with positive results. However, there is still seldom information on the type of rTMS protocols that deliver the best clinical improvement in AD.

**Objetive:** To compare the clinical response between a simple stimulation protocol on the left dorsolateral prefrontal cortex (lDLPFC) against a complex protocol using six regions of interest.

**Methods:** 19 participants were randomized to receive any of the protocols. The analysis of repeated measures evaluated the change.

**Results:** Both protocols were equally proficient at improving cognitive function, behavior and functionality after 3 weeks of treatment, and the effects were maintained for 4 weeks more without treatment.

**Conclusion:** We suggest rTMS on the lDLPFC could be enough to provide a clinical response, and the underlying mechanisms should be studied.

## Introduction

Alzheimer’s disease (AD) is a neurodegenerative disorder of uncertain etiology that affects around 46.8 million people worldwide. It is characterized by progressive cognitive decline that affects behavior and function. In AD, there is a limited pharmacological treatment efficacy (1). Clinical trials using rTMS in AD have found positive effects on cognition, behavior and function (2-4). High frequency (HF) rTMS applied on the right or left DLPFC has shown improvement in language abilities, in cognitive function, functionality and behavior for up to 3 months (5). Other studies have shown similar results on cognition, behavior and function using HF applied over several cortical sites in a complex design (6). This complex design may include at least 6 cortical targets, and it is suggested to be more beneficial than DLPFC (7,8). In this study we compare two rTMS modalities: simple stimulation of the lDLPFC vs. complex stimulation of 6 regions related to AD's known brain affected areas.

## Materials and Methods

The study was conducted according to the Declaration of Helsinki and was approved by the ethics committee of the Instituto Nacional de Psiquiatría (No. CEI/C/049/2015). The study was registered in ClinicalTrials Gov. Sample size was calculated using a repeated measures design with an alpha of 0.05, a power of 90% and effect size of 0.5 (9), for a sample size of 22 participants (11 per group). We recruited 33 patients with diagnosis of dementia, and 19 with diagnosis of AD were included in the clinical trial (criteria in Supplementary Methods).

The design was longitudinal, single blind and the patients were randomized for the treatment groups: 1) lDLPFC, and 2) 6-ROIs. In both groups we used rTMS for 3 weeks where each session consisted of 30 trains of 10 sec of duration separated by 1 min rests, with a frequency of 5 Hz at 100% of the motor threshold. The lDLPFC arm was applied using a MagPro R30 (Magventure, Denmark) with an 8-shape coil model MCF-B70 and consisted of stimulation only in the lDLPFC, localized using the 10-20 system, 1,500 pulses per session. The 6-ROIs arm was applied using a MagPro (Dantec, Denmark) with an 8-shape coil model MC-B70 and consisted of stimulation in 6 different regions with the following protocol. For this, we alternated stimulation between regions each day; on one day we stimulated three areas: a) Broca’s area, b) Wernicke’s area and c) lDLPFC, and the next day we stimulated three areas: d) lpSAC, e) rpSAC, f) rDLPFC, both days with 500 pulse per area, for 1,500 pulses per session.

Our primary outcome measure was changes in cognitive function, using ADAS-cog at baseline, week 3 (during stimulation), and week 7 (4 weeks after the last rTMS session). Our secondary outcome measures were changes in: cognitive measures (Mini Mental State or MMSE), behavioral symptoms report (NPI), depression symptoms (GDS-Yesavage), functionality evaluation (IDDD) and Clinical Global Impression (CGI). Clinical improvement in cognition was defined as: 1) reduction of 4 points in ADAS-cog and, 2) an increase of 1.5 - 2 higher than baseline MMSE. In the other variables, any positive change in comparison to the baseline was considered improvement. We performed descriptive and inference analysis of all variables using IBM SPSS ver. 22 (International Business Machines Corp, New York, USA). For outcome measures analyses we used Repeated-measures ANOVA for within-subject factors (Baseline, Week3 and Week7) and particularly for Mini Mental there were four measures (Baseline, Week1, Week2, Week3 and Week7), with an alpha of 0.05.

## Results

For the lDLPFC group we included 10 patients (6 female, mean age 73.30 (± 6.03)), and for the 6-ROIS group we included 9 patients (5 female, mean age 71 (± 4.27)). We had no drop-outs in our study and four patients merely reported a transitory mild headache after rTMS as a secondary effect. The baseline scores were similar in both groups in all measurements. We found significant clinical improvement in primary and secondary outcome measures after rTMS treatment (Week3) in both groups. We found that the patients maintained this effect at follow-up after one week without treatment (Week7). We did not find significant differences between groups in any measure, meaning that both rTMS modalities were equally beneficial (Figure 1).

## Discussion

In our study we compared two different rTMS modalities of treatment in AD patients. We found that both modalities improved the patients’ cognitive, behavioral and functional measures equally. Therefore, the benefit of rTMS in AD may mostly rely on lDLPFC stimulation The neurophysiological mechanisms related to the beneficial effects of rTMS are still poorly understood. A proposed mechanism suggests an increase in processing efficiency due to the direct modulation of cortical areas or networks by adjusting pathological brain patterns of activity and inducing improved activity patterns. The absence of a difference between rTMS modalities may be explained by the stimulation of the lDLPFC in both. The lDLPFC may act as an important hub for network integration in cognition and behavior, which is pathologically disrupted in AD. Therefore, the use of rTMS in this region may improve the network activity and integration, which may be directly related to clinical improvement (10).

A limitation of our study was the lack of a sham group. However, the goal of our study was to directly compare two different modalities, complex vs. simple rTMS and not to asses the already demonstrated clinical effect of in both modalities (8). We decided to first investigate both modalities directly as finding differences between modalities may have justified a sham clinical trial. Another limitation is the lack of a neuronavigator to determine the precise localization of each stimulated region. Nevertheless, the 10-20 system is the most used in the daily clinical practice due, hence, results may be more relevant in that regard. In summary, we found that rTMS of the lDLPFC and the stimulation of six different regions was equally beneficial for AD patients at the safe frequency of 5 Hz. Therefore, we do not see a benefit from applying the complex stimulation.

## Acknowledgements

We would like to thanks Dr. Carlos H. Berlanga Cisneros for his supervision in this project. The project was funded by the Instituto Nacional de Psiquiatría "Ramón de la Fuente Muñiz” No. SIC-16-001.

## Figure Legends

**Figure 1.** a) Study flowchart. b) The study design consisted on an active stimulation period for 3 weeks and a passive period without stimulation of 4 weeks. c) ADAS-Cog, d) MMSE, e) IDDD, and f) NPI line plots with error bars showing standard deviation and results from the within subjects repeated measures ANOVA. lDLPFC = Simple stimulation group on the left dorsolateral prefrontal cortex; 6-ROIs = Complex stimulation group on 6 regions of interest.; ALL = All clinical questionnaires (ADAS-cog, MMSE, NPI, IDDD, GDS-Yesavage and CGI); MMSE = Mini Mental State; ADAS-Cog = Alzheimer’s Disease Assessment Scale – Cognitive; NPI = Neuropsychiatric Inventory Cummings; IDDD = Interview for Deterioration in Daily Living Activities in Dementia; GDS-Yesavage = Yesavage Geriatric Depresion Scale; CGI = Clinical Global Impression; BL = Baseline; W = Week; F = f-ratio; p = p-value.

